# Short cracks in knee meniscus tissue cause strain concentrations, but do not reduce ultimate stress, in single-cycle uniaxial tension

**DOI:** 10.1101/292623

**Authors:** John M. Peloquin, Michael H. Santare, Dawn M. Elliott

## Abstract

Tears are central to knee meniscus pathology and, from a mechanical perspective, are crack-like defects (cracks). In many materials, cracks create stress concentrations that cause progressive local rupture and reduce effective strength. It is currently unknown if cracks in meniscus have these consequences; if they do, this would have repercussions for management of meniscus pathology. The objective of this study was to determine if a short crack in meniscus tissue, which mimics a preclinical meniscus tear, (a) causes crack growth and reduces effective strength, (b) creates a near-tip strain concentration, and (c) creates unloaded regions on either side of the crack. Specimens with and without cracks were tested in uniaxial tension and compared in terms of macroscopic stress–strain curves and digital image correlation strain fields. The strain fields were used as an indicator of stress concentrations and unloaded regions. Effective strength was found to be insensitive to the presence of a crack (potential effect < 0.86 s.d.; β = 0.2), but significant strain concentrations, which have the potential to lead to long-term accumulation of tissue or cell damage, were observed near the crack tip.

## 1 Introduction

Tears are the most common pathology to afflict the knee meniscus. Their direct symptoms include pain, mechanical deficiency, and increased risk of osteoarthritis (Englund et al. 2012; Øiestad et al. 2009; Cohen et al. 2007; Englund et al. 2008). Meniscus tears can also cause serious secondary harm, as the most straightforward treatment for a tear—partial or total meniscectomy—further increases the risk of osteoarthritis (Magnussen et al. 2009; Shelbourne and Gray 2000; Feucht et al. 2015; Anetzberger et al. 2014; Petty and Lubowitz 2011). Furthermore, repair of a torn meniscus, especially when the tear is in the inner avascular region, is not always feasible (Arnoczky and Warren 1983; Englund et al. 2012). Some in vivo meniscus tears occur suddenly, suggestive of immediate rupture from a single overload event (Fink et al. 2018; Greis et al. 2002; Wagemakers et al. 2008; Cox et al. 2009; Lento and Akuthota 2000; Poulsen and Johnson 2011; Paul et al. 2003), whereas others grow over time (Kumm et al. 2015; Khan et al. 2016). Although short (< 5–10 mm) tears often do not merit immediate clinical concern (Lento and Akuthota 2000; Stärke et al. 2009; Roeddecker et al. 1994; Lento and Akuthota 2000; Seil et al. 2009), they have the potential to grow and become more serious with time. To aid efforts to prevent and to repair meniscus tears, it is important to understand the mechanics of how they form and grow.

A meniscus tear, viewed from an engineering perspective, is a macroscopic crack-like defect. In this work, “meniscus tear” will be used to refer to the clinical pathology, and “meniscus crack” will be used in the engineering sense to mean a macroscopic crack-like defect in the meniscus, regardless of whether it is a natural meniscus tear or an artificial crack-like defect such as a cut. A crack typically creates a stress concentration at its tip, which often, but not always, causes crack growth and reduces ultimate stress (maximum engineering stress prior to rupture) (Janssen et al. 2002; Anderson 2005; Broberg 1999). Study of this type of failure comprises the field of fracture mechanics. The central question concerning meniscus cracks is: do cracks reduce the stress-bearing capacity of meniscus tissue, or is meniscus tissue insensitive to them?

The effect of a crack depends on the material’s fracture toughness and the severity of the crack-induced stress concentration (Figure 1) (Anderson 2005; Taylor et al. 2012; Janssen et al. 2002). If the stress concentration is large relative to the material’s fracture toughness, it causes local rupture at the crack tip, crack growth, and a reduction in ultimate stress (fracture mechanics region, Figure 1). In this case fracture mechanics approaches should be used to quantify rupture. If the stress concentration is small relative to the material’s fracture toughness, the crack has negligible effect, and the ultimate stress is equal to the ultimate tensile strength (limit stress analysis region, Figure 1). In this case, limit stress analysis (comparison of applied stress to material properties, such as ultimate tensile strength) should be used to quantify rupture. Demonstration of limit stress analysis-dominant rupture would not necessarily mean that defects such as microcracks are uninvolved in failure, but it would indicate that the primary impetus of rupture was *not* macroscale crack growth. To permit accurate failure analysis of meniscus bearing a short tear, it is crucial to determine if meniscus tissue containing a short crack ruptures by fracture mechanics-dominant processes or limit stress analysis-dominant processes.

**Figure 1:**
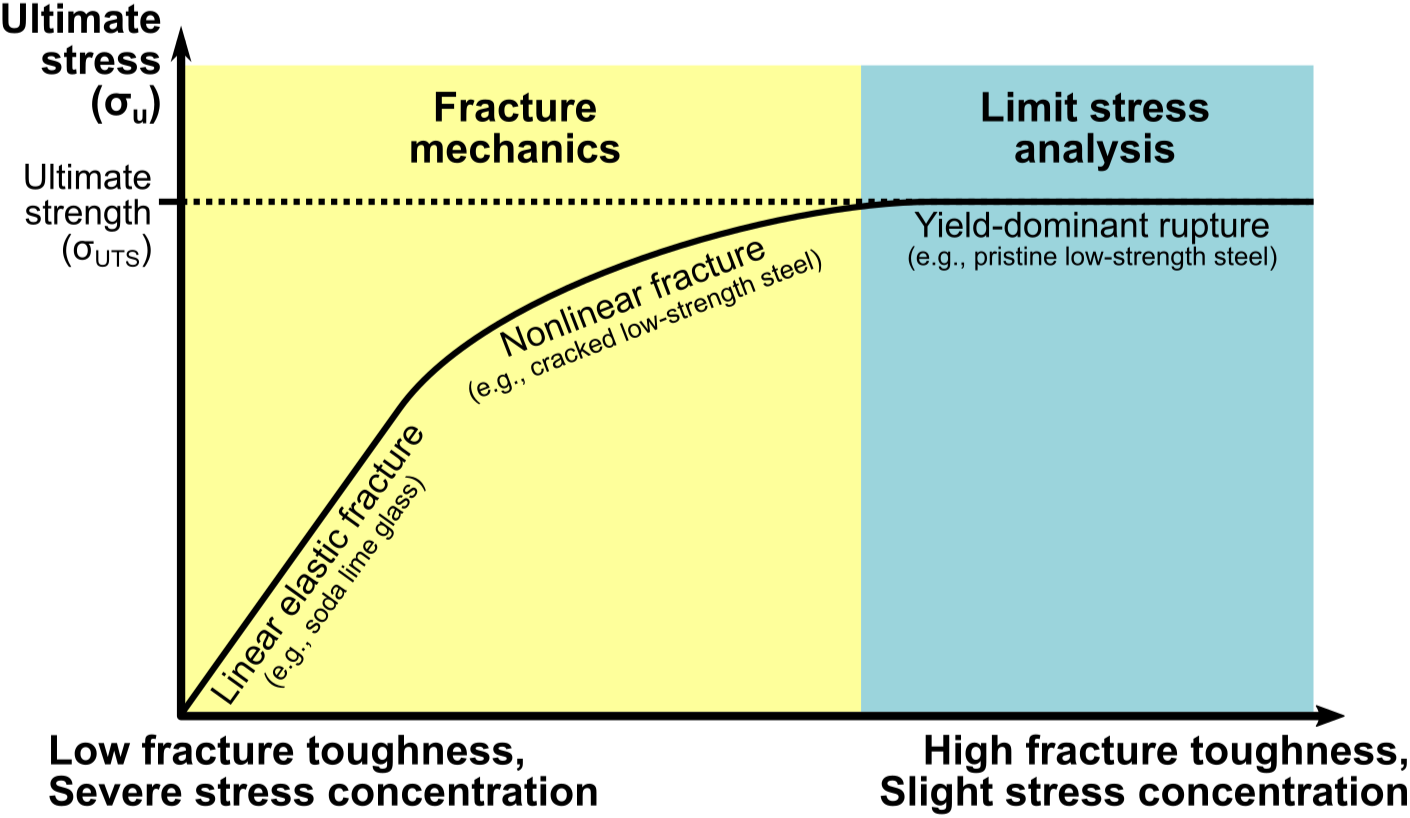
Schematic of ultimate stress variation with fracture toughness and stress concentration severity. Fracture mechanics governs failure when a local stress concentration is large relative to the material’s fracture toughness. Limit stress analysis governs failure when the material is sufficiently tough to tolerate any stress concentrations that are present. In the fracture mechanics case, σ_u_ < σ_UTS_; in the limit stress case, σ_u_ = σ_UTS_ (Anderson 2005; Taylor et al. 2012; Janssen et al. 2002).

The idea that the meniscus ruptures by fracture mechanics-dominant processes has some experimental support. Meniscus tears grow in vivo (Kumm et al. 2015; Khan et al. 2016), suggesting that crack growth in the fracture mechanics sense may have a role. Superficial zone articular cartilage, which is similar to meniscus, has been observed to rupture by fracture mechanics-dominant processes (Chin-Purcell and Lewis 1996; Taylor et al. 2012). However, cracks in tendon, which like meniscus contains highly aligned collagen fascicles, tend to become blunt instead of growing (Schechtman and Bader 1997; Von Forell and Bowden 2014b; Von Forell and Bowden 2014a; Andarawis-Puri et al. 2009; Locke et al. 2017; Szczesny et al. 2015). These findings suggest that limit stress analysis is more appropriate for cracked tendon, and by extension possibly also for meniscus. It remains uncertain which rupture process is relevant for meniscus, and mechanical tests are needed to resolve this uncertainty.

To determine if cracked meniscus fails by fracture mechanics-dominant processes, we chose to compare the ultimate stress of cracked meniscus specimens to identically tested intact control specimens. This test has been specifically recommended for application to fibrous soft tissues for which the dominant failure process is not known (Taylor et al. 2012). Cracked specimens were prepared with crack length much longer than any pre-existing defects that may have been present. Because stress concentration severity increases with crack length, cracked specimens will lie to the left of the control specimens on Figure 1. If the cracked specimens lie in the fracture mechanics region, they will have ultimate stress less than controls. If the cracked specimens instead lie in the limit stress analysis region, they will have ultimate stress equal to the controls. Observation of either outcome is valuable; either the results provide a necessary foundation to support subsequent fracture toughness measurements, or they rule out risk of crack-induced weakness for the tested crack geometry and loading conditions.

Independent of which process governs rupture, the potential for cracks to alter stress and strain field within the meniscus is also of interest—specifically, the potential for near-tip stress concentrations and unloaded regions on either side of the crack. A stress and strain concentration may cause stress to locally exceed the elastic limit and hence cause local tissue or cell damage. Conversely, unloaded regions may cause long-term cell-mediated degeneration by abolishing mechanical signals necessary for homeostasis. The idea that a near-tip stress concentration produces local damage is supported by the observation that tissue near the tip of a naturally occurring meniscus tear takes less energy to rupture than tissue in a crack-free contralateral control meniscus (Roeddecker et al. 1994). Concerning unloaded regions, a crack in a meniscus-mimicking synthetic scaffold has been observed to reduce strain in the regions that include material cut by the crack (Bansal et al. 2017; Tsinman et al. 2017). Quantification of any crack-induced stress–strain concentrations and unloaded regions is needed to determine if these mechanical phenomena may have a role in meniscus tear pathology.

The particular objectives of this study were, for meniscus tissue with a short crack: (a) to determine whether fracture mechanics or limit stress analysis is appropriate for failure analysis; (b) to quantify strain concentrations in the region near the crack tip, and (c) to quantify the degree of unloading in the cut material to either side of a crack. Meniscus specimens with and without artificially introduced cracks were tested in both cir-cumferential and radial uniaxial tension so as to examine both fiber and matrix rupture phenomena, and because tensile load support is a key aspect of meniscus function (Kawamura et al. 2003; Freutel et al. 2014; Jones et al. 1996; Kolaczek et al. 2016; Upton et al. 2006; Spilker et al. 1992). To test the hypothesis that meniscus cracks reduce ultimate stress and cause rupture by fracture mechanics-dominant processes (objective a), ultimate stress was compared between cracked and crack-free specimens as recommended by Taylor et al. (2012). To test the hypothesis that near-tip strain concentrations are present and to measure their magnitude (objective b), strain fields were measured using digital image correlation (DIC) and strain in cracked specimens was compared between their near-tip and away-from-tip regions. Similarly, to test the hypothesis that the regions to either side of a crack, which contain cut material, are unloaded (objective c), strain in cracked specimens was compared between their cut and intact regions. Results are discussed in terms of their significance with respect to whole-meniscus damage and failure.

## 2 Methods

### 2.1 Specimen preparation

Specimens were prepared for three test groups—circumferential tension with a 90° edge crack, circumferential tension with a 45° center crack, and radial tension with a 90° edge crack—with corresponding circumferential and radial tension crack-free control groups (Figure 2). As established by prior work (Peloquin et al. 2016), circumferential specimens were made with expanded grip regions to facilitate fiber loading. The specimens were prepared from adult (age > 30 months) bovine menisci.

**Figure 2:**
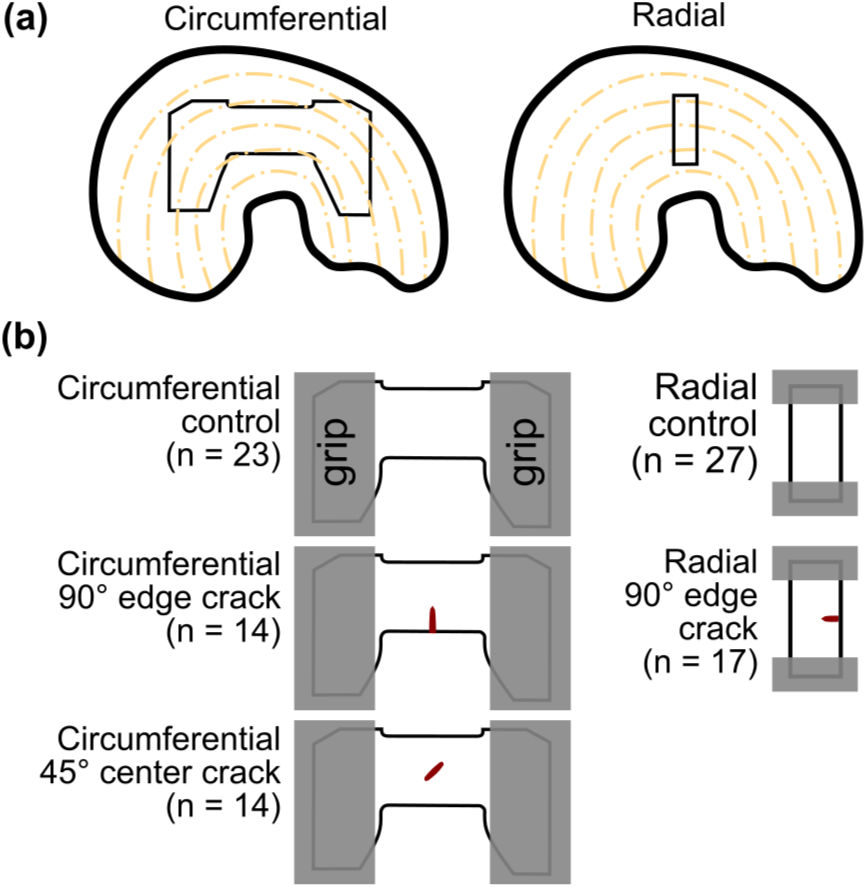
Specimen schematics. (a) Sites of specimen dissection. (b) Specimen shapes for each of the analysis groups (circumferential tension 90° edge crack, circumferential tension 45° center crack, and radial tension 90° center crack) and their crack-free controls.

Menisci (mixed lateral & medial) were purchased from Animal Technologies, Inc. (Tyler, TX) and stored frozen (−20 °C). One specimen per meniscus was prepared using a Leica SM2400 sledge microtome, with a freezing stage, to plane the meniscus to a target thickness of 1.0–1.5 mm, with the specimen plane normal to the proximal–distal axis (Lechner et al. 2000). The specimen was then trimmed to its final in-plane dimensions, its cross-sectional area measured by scanning across its width with a laser displacement sensor (Szczesny et al. 2012; Favata 2006), and its surface airbrushed with Verhoeff’s stain to facilitate digital image correlation from test video. Each edge crack was created by incision with a fresh #11 scalpel blade to a target length of 3 mm. Each center crack was created by incision with a fresh razor blade broken to the target crack length of 3 mm. Measured specimen dimensions are reported in Table 1. Except when the specimens were being actively manipulated, they were kept covered under PBS-dampened gauze to minimize dehydration.

**Table 1:** Specimen dimensions (mean ± sd).

| Group | Width (mm) | Grip-to-grip length (mm) | Thickness (mm) | Crack length (mm) |
| --- | --- | --- | --- | --- |
| Circumferential control | $8.0 \pm 2.1$ | $16.2 \pm 2.3$ | $1.2 \pm 0.2$ | |
| Circumferential 90° edge crack | $9.2 \pm 2.4$ | $16.5 \pm 1.9$ | $1.0 \pm 0.2$ | $2.9 \pm 0.4$ |
| Circumferential 45° center crack | $9.8 \pm 2.0$ | $16.0 \pm 2.1$ | $1.2 \pm 0.3$ | $3.1 \pm 0.3$ |
| Radial control | $5.6 \pm 1.0$ | $8.1 \pm 1.6$ | $1.5 \pm 0.3$ | |
| Radial 90° edge crack | $5.9 \pm 1.6$ | $8.1 \pm 1.8$ | $1.5 \pm 0.3$ | $2.0 \pm 0.4$ |

### 2.2 Tensile test protocol

Tensile testing was done following standard procedures Peloquin et al. (2016), LeRoux and Setton (2002), Proctor et al. (1989), Anderson et al. (1993), Tanaka et al. (2014), Sweigart and Athanasiou (2005), Tissakht and Ahmed (1995), and Lechner et al. (2000). The tensile test protocol consisted of (a) 20 kPa preload, which was used to establish the specimen’s undeformed reference length, (b) 10 cycles of preconditioning to 4% grip-to-grip engineering strain, and (c) stretch to failure. The displacement rate was 0.5 mm/s. Video of the test was recorded at 15 fps and 1392 × 1000 px. The image scale was ~ 40 px/mm for circumferential specimens and ~ 60 px/mm for radial specimens.

Ruptures of tested specimens were classified according to published definitions (Peloquin et al. 2016). Tests with rupture or slip inside the grips (longitudinal split, gripped region failure, or no rupture) were excluded from analysis because mean stress in the failure region cannot be accurately calculated for these failure types. The counts of each type of rupture are presented in Table 2. The counts of tests used for analysis are given by group in Figure 2b. The rupture types that were used for analysis do not systematically differ in their stress–strain response under this protocol (Peloquin et al. 2016).

**Table 2:** Counts for each type of rupture by specimen group.

| Group | Rupture Type | Included in analysis | Control | 90° edge crack | 45° center crack |
| --- | --- | --- | --- | --- | --- |
| Circumferential | Midsubstance rupture | Yes | 5 | 2 | 4 |
|  | Mixed rupture | Yes | 11 | 12 | 8 |
|  | Grip line rupture | Yes | 7 | 0 | 2 |
|  | Gripped region failure | No | 8 | 6 | 5 |
|  | Longitudinal split | No | 0 | 2 | 0 |
|  | No rupture | No | 5 | 0 | 2 |
| Radial | Midsubstance rupture | Yes | 9 | 11 |  |
|  | Mixed rupture | Yes | 3 | 3 |  |
|  | Grip line rupture | Yes | 15 | 3 |  |
|  | Gripped region failure | No | 0 | 1 |  |
|  | Longitudinal split | No | 0 | 0 |  |
|  | No rupture | No | 1 | 2 |  |

### 2.3 Stress and strain data processing

The tensile tests’ stress (σ)–strain (ε) curves were summarized using ultimate stress σ_u_, strain at ultimate stress ε_u_, tangent modulus (measured at the yield point), yield stress, and yield strain. Strain was calculated using the current grip-to-grip length *l* and the reference grip-to-grip length *l*_0_ and reported as stretch ratio (λ = *l*/*l*_0_) or Lagrange strain (*ϵ* = 1/2[λ^2^ − 1]). The yield point was identified as the inflection point in the stress–strain curve as in Peloquin et al. (2016). Note that for tissue tests the yield point is not customarily equated with the elastic limit, in part because the role of plasticity with respect to tissue remains ambiguous (Peloquin et al. 2016; Espejo-Baena et al. 2006; Veres et al. 2013; Danso et al. 2014; Jones et al. 2014; Palmer et al. 2009; Smith et al. 2008; Haslach et al. 2018). For cracked specimens, the uncut transverse cross-sectional area (thickness × (total width – transverse axis component of crack length)) was used for stress calculations.

Lagrange strain fields were computed across the specimen surface from the video recordings using 2D DIC (Vic-2D 2009; Correlated Solutions, Columbia, SC) (Andarawis-Puri et al. 2009; Mallett and Arruda 2017; Peloquin et al. 2016). Three DIC strain components were analyzed: longitudinal strain (grip-to-grip direction; E_xx_), transverse strain (E_yy_), and shear strain magnitude (|E_xy_|). The correlation window (subset size) was 0.7 × 0.7 mm, and incremental correlation, exhaustive search, and low pass filtering were enabled. Strain accumulation error from incremental correlation was negligible; for this test apparatus and the number of frames analyzed, strain error for each pixel is ≤ 0.001. Post-processing used a 15 px exponential decay filter. Trial runs during study development showed that only the subset size setting had a strong influence on the calculated strain field. The choice of 0.7 × 0.7 mm subset size was a compromise between the needs of (a) minimizing spurious correlations, which favors a large correlation window, and (b) retaining spatial resolution to identify strain concentrations and inter-fascicle sliding, which favors a small correlation window.

Quantitative analysis of strain fields was done by comparison of strain statistics between regions of interest (ROIs). To test for the existence of strain concentrations near the crack tip, the DIC strain field was partitioned into near-tip and away-from-tip ROIs (Figure 5a). The ROI size was chosen to be of similar size to the strain concentrations visible in the strain field images. The near-tip ROIs for analysis of E_xx_ extended, relative to the crack tip, +0.7 mm in the crack pointing direction and ±0.7 mm in the loading direction. The near-tip ROI size was set to approximately the subset window size because this is the smallest size that is reasonable for analyzing spatial variation. The near-tip ROIs for analysis of E_yy_ and E_xy_ were similar defined to extend +0.7 mm in the crack pointing direction and, because features in the E_yy_ and E_xy_ strain fields were elongated in the loading direction, ±1.05 mm in the loading direction. To test for differences in strain between tissue cut and not cut by a crack, “cut” and “intact” ROIs were also defined (Figure 7a). Comparisons of DIC strain statistics between paired (a) near-tip and away-from-tip and (b) cut and intact ROIs were made at stress levels from 0.25σ_u_ to 0.9σ_u_ for circumferential specimens and, due to loss of DIC tracking at high strain for several radial specimens, 0.25σ_u_ to 0.7σ_u_ for radial specimens. Measurement at relative stress levels ensures that each measurement reflects a similar phase of the stress–strain curve across all specimens.

### 2.4 Statistics and inference

The hypothesis that cracks reduce the ultimate stress of meniscus (related to objective a) was tested using unpaired Welch t-tests to compare ultimate stress between cracked and control specimens for each of the three test cases (circumferential tension 90° edge cracks, circumferential tension 45° center cracks, and radial tension 90° edge cracks). Since this hypothesis was specifically that cracked σ_u_ < control σ_u_, comparisons were made using a one-sided t-test to decrease the chance of type II error. The other stress–strain parameters did not have a specific hypothesized direction of change and so were compared between cracked and control specimens with two-sided t-tests. The type I error rate for all statistical tests was set to 0.05.

The hypothesis that cracks generate strain concentrations near their tips was tested by comparing DIC strain (objective b) in the near-tip ROI to the away-from-tip ROI (Figure 5a). Similarly, the hypothesis that cracks reduce strain in the region containing cut material (objective c) was tested by comparing DIC strain in the cut ROI to the intact ROI (Figure 7a). The between-ROI differences in DIC strain as t-tests had skewed and very heavy-tailed distributions, making t-tests unsuitable. Therefore comparisons of DIC strain between ROIs within each specimen were done using the mean strain in each ROI by paired Wilcoxon tests.

## 3 Results

### 3.1 Effect of cracks on stress–strain curve parameters

Whether fracture mechanics or limit stress analysis is appropriate for failure analysis of cracked meniscus tissue was determined by comparing ultimate stress (σ_u_) for each group of cracked specimens with its corresponding control group (Figure 3; see Figure 1 for rationale). None of the crack groups (circumferential 90° edge crack, circumferential 45° center crack, and radial 90° edge crack; Figure 2) had ultimate stress significantly less than their corresponding controls (p > 0.2 for all). The study had power = 0.8 to detect a 0.86 s.d. decrease in circumferential ultimate stress and a 0.78 s.d. decrease in radial ultimate stress, so this negative result indicates that there was no meaningful difference. Therefore the specimens failed by limit stress dominant processes.

**Figure 3:**
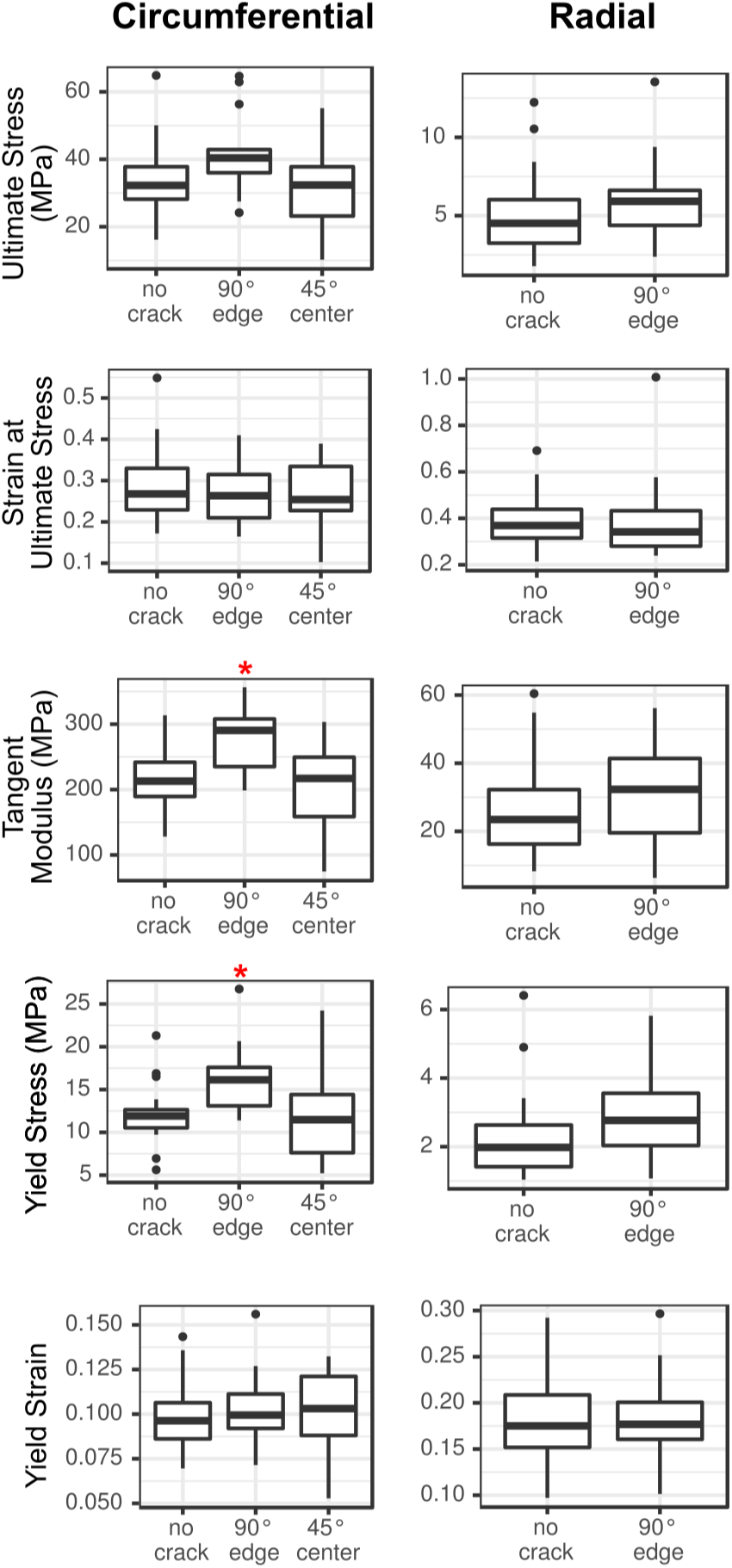
Comparison of stress–strain curve parameters between cracked specimens and their corresponding controls. The boxplots are in Tukey’s style. * indicates cracked case ≠ control case, Welch t-test, p < 0.05.

The other stress–strain curve parameters—strain at ultimate stress, tangent modulus, yield stress, and yield strain—were also compared between the crack groups and their controls (Figure 3). The full table of stress–strain curve statistics is provided in supplemental information (Table S1), and the stress–strain curves are plotted in Figure 4 for qualitative comparison. Tangent modulus and yield stress were significantly greater in circumferential 90° edge crack specimens by 1.5–6.8 MPa and 18–88 MPa respectively (95% CIs). No other stress–strain parameter comparison showed a significant difference between the crack and control groups.

**Figure 4:**
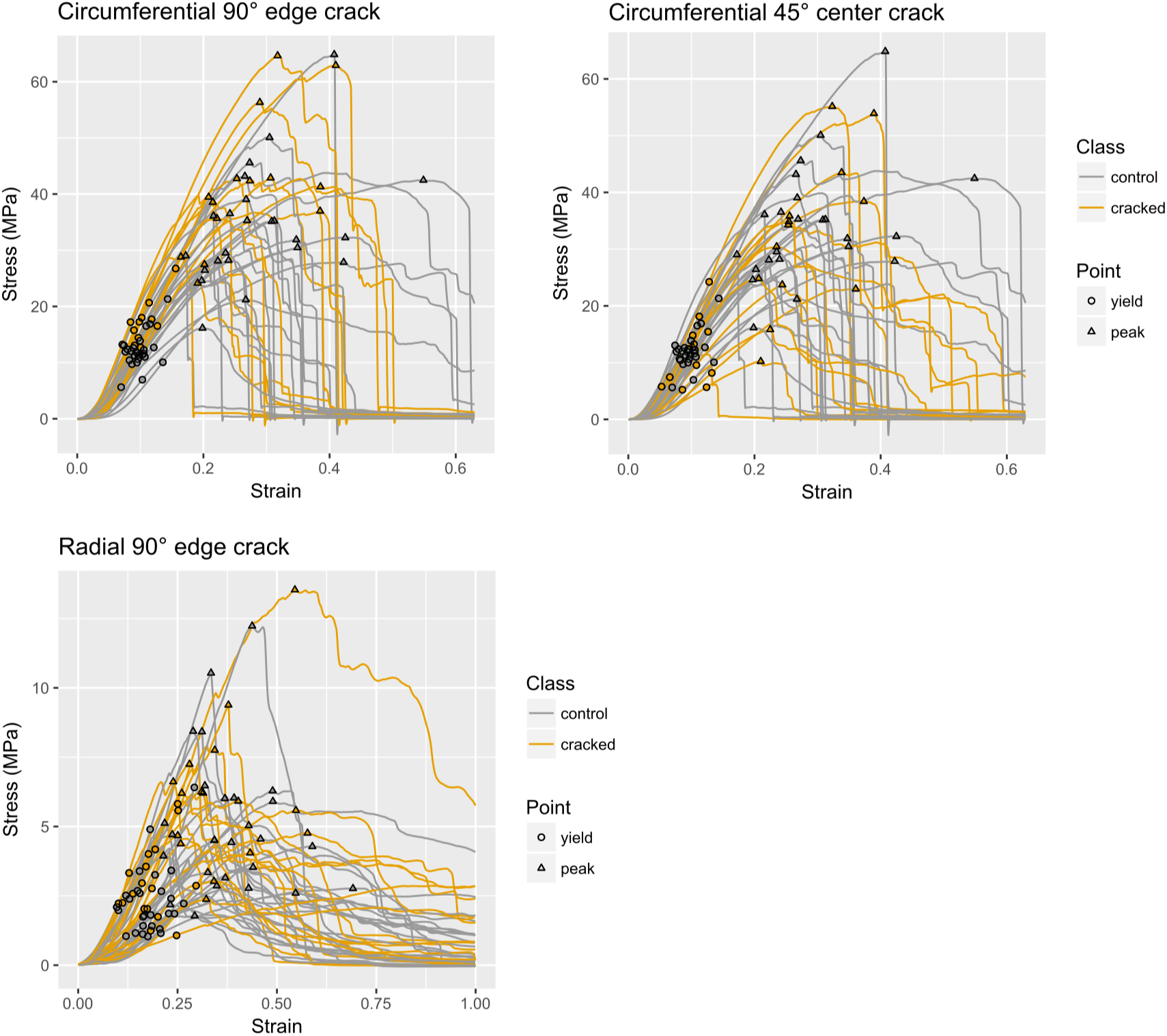
Stress-strain curves for all analyzed specimens.

### 3.2 Effect of cracks on strain fields

The presence of crack-induced strain concentrations was tested by comparing, for each specimen, mean DIC strain near and away from the crack tip. Longitudinal strain (E_xx_) was significantly greater near the crack tip for all groups at every stress level examined (Figure 5). Transverse strain (E_yy_) was not significantly different near the crack tip in circumferential tension specimens, but for σ ≥ 0.5σ_u_ was significantly more compressive near the tip in radial edge crack specimens. Shear strain magnitude (|E_xy_|) was significantly greater near the crack tip in circumferential edge crack specimens for σ ≥ 0.8σ_u_ and in circumferential center crack and radial edge crack specimens for all stress levels examined. The complete table of strain field statistics is provided in supplemental information (Table S2). These results demonstrate that cracks in meniscus tissue create strain concentrations at their tips.

**Figure 5:**
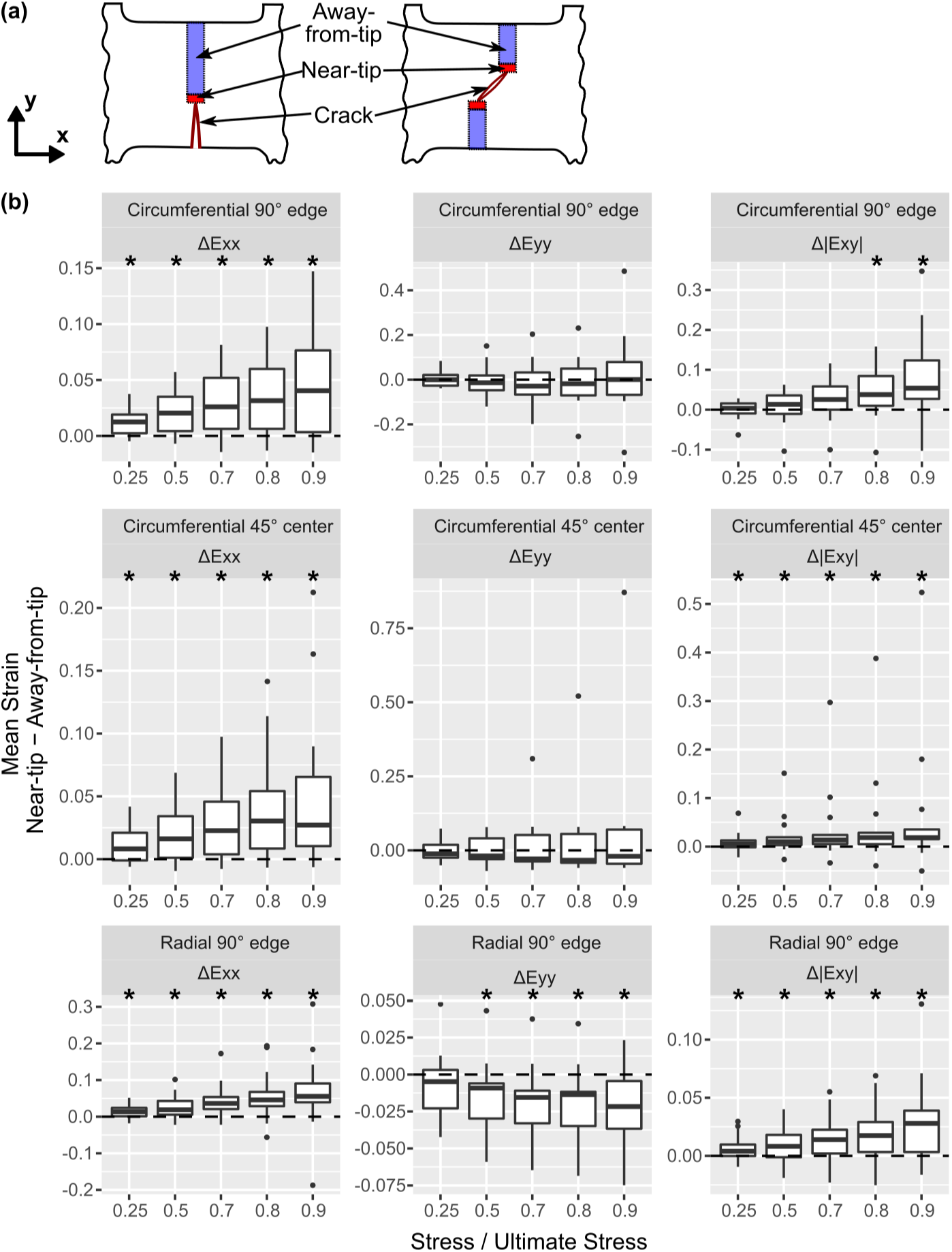
Strain field differences between near-tip and away-from tip ROIs within specimens. (a) Schematic of ROI definitions. (b) Tukey boxplots. * indicates near ≠ away, Wilcoxon test, p < 0.05.

The strain fields were qualitatively examined in terms of the near-tip strain concentrations’ size and shape, as well as the strain fields’ overall appearance. Strain fields for three representative cracked specimens are shown in Figure 6. Strain fields for control specimens were similar to those in Peloquin et al. (2016). The mottled appearance of each test’s strain field, specifically the pattern of blobs of positive and negative strain, was established early in the test (~ 0.01 strain) and retained the same pattern through failure.

**Figure 6:**
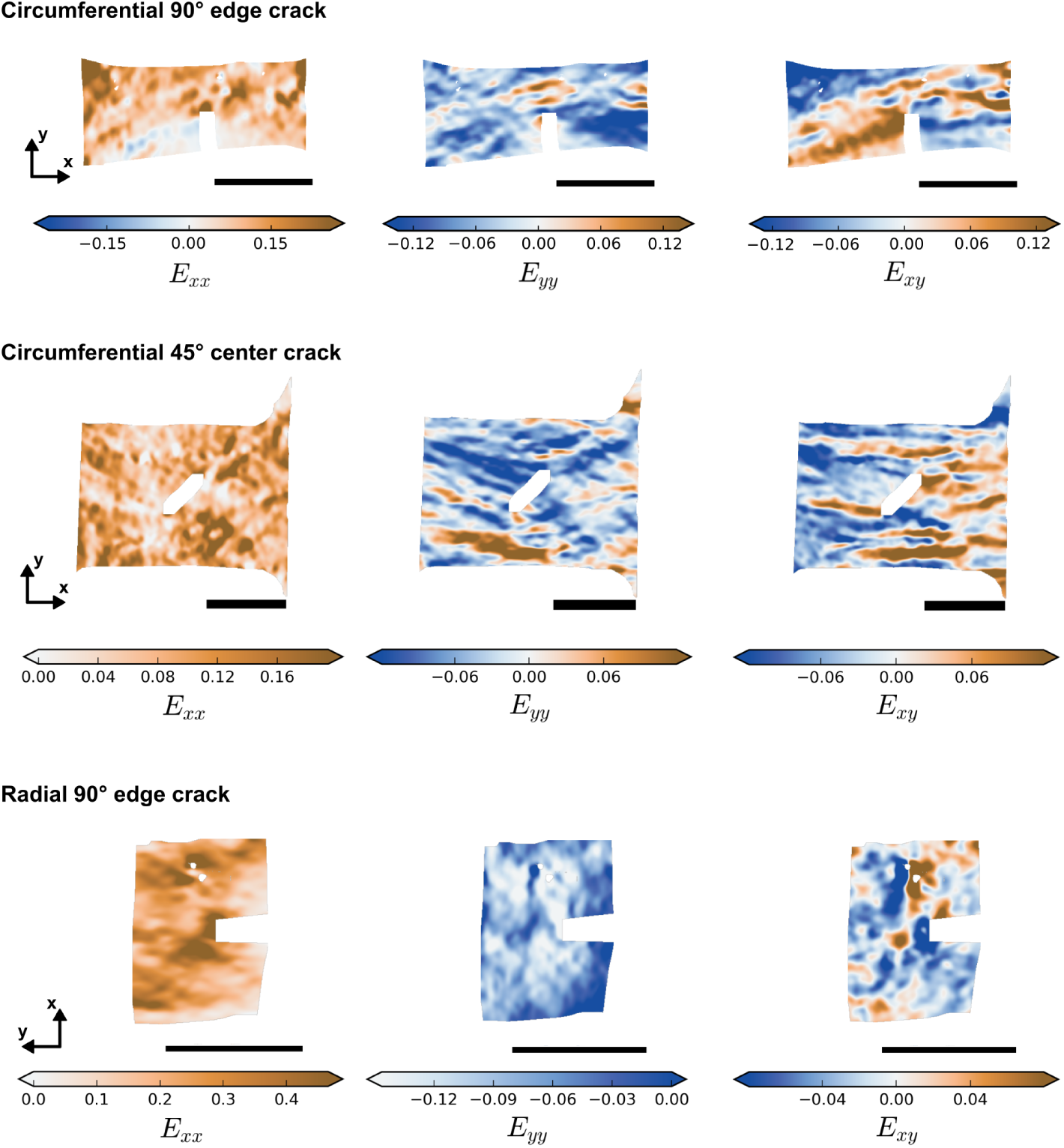
Representative strain fields for cracked specimens in each group at σ = 0.7σ_u_. Scale bar = 5 mm.

Although the strain fields showed great variation between specimens, they had some consistent features. The near-tip E_xx_ concentrations in circumferential and radial specimens tended to be small and subtle, extending ≈ 0.5 mm from the crack. In radial specimens, the near-tip E_xx_ concentrations sometimes (5/17 tests) grew to extend across the specimen as σ approached σ_u_ and as rupture progressed from the crack tip. Near-tip E_xy_ concentrations in both circumferential and radial specimens took the form of bands that extended longitudinally across 50–100% of the specimen. The crack-associated E_xy_ bands switched sign between sides of the crack. For all strain components, strong strain concentrations tended to develop at multiple places within each specimen. E_xx_ and E_yy_ were on average greater near the crack tip because a strain concentration occurred there consistently across specimens, not because it was uniquely strong within each specimen. Exy concentrations, however, were more specific to the presence of a crack; in ~ 1/3 of circumferential tests and ~ 1/2 of radial tests, the strongest and most extensive shear band was crack-associated.

To test if the cracks created unloaded regions in cracked specimens, mean strain in the cut ROI (on the crack flanks) was compared to the intact ROI (Figure 7; also plotted as an attenuation ratio in Figure S1). Mean E_xx_ in the cut ROI was significantly less than in the intact ROI for all crack groups at all stress levels. In edge crack specimens, both circumferential and radial, this effect almost completely abolished E_xx_ in the cut ROI. Mean E_yy_ in the cut ROI in circumferential specimens was not significantly different from the intact ROI, but in radial specimens it was significantly greater than in the intact ROI. Mean |E_xy_| in the cut ROI of circumferential edge and center crack specimens was less than in the intact ROI, whereas in radial specimens it was greater. As E_xx_ is the strain component most directly related to longitudinal stress, the large reductions in the cut ROIs’ mean E_xx_ values indicate that the crack did unload the cut ROI.

**Figure 7:**
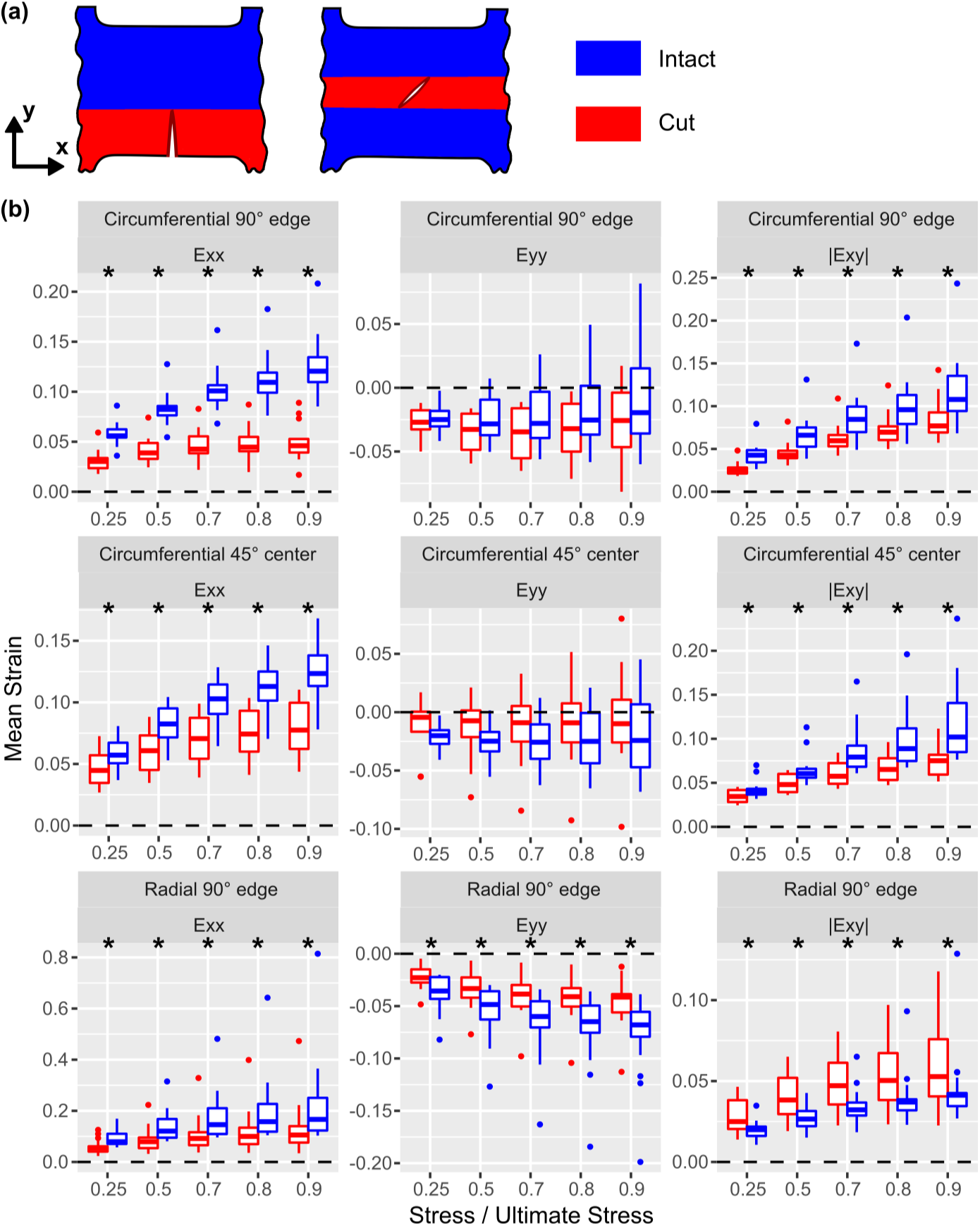
Strain field differences between cut and intact ROIs within specimens. (a) ROI definitions. (b) Tukey boxplots. * indicates cut ≠ intact, paired Wilcoxon test, p < 0.05.

### 3.3 Rupture morphology

Rupture morphology of cracked specimens, particularly whether rupture occurred by crack growth, was summarized to support interpretation regarding rupture mechanisms. Rupture in the circumferential edge crack specimens proceeded predominantly by inter-fascicle sliding, creating broad, irregular ruptures with extensive interdigitating fiber pullout (Figure 8a). In 13 out of 14 circumferential edge crack tests, the region near the crack tip was the first to display visible rupture, but the crack blunted rapidly by inter-fascicle shear (Figure 8b) and final rupture occurred simultaneously over a broad region. Crack blunting by inter-fascicle shear began very early in the test, between 0.2σ_u_ and 0.5σ_u_. Rupture in circumferential center crack specimens occurred similarly, except that the ruptures at the crack tip were smaller and developed in parallel with separate sites of rupture at the grip line (5 tests) or specimen edge (9 tests). These separate sites of rupture typically joined the crack-associated rupture as they grew (10 tests; Figure 8c). Inter-fascicle shear was ubiquitous in both edge and center crack circumferential specimens, and often caused the rupture line to zigzag across the specimen between zones of interdigitating fiber pullout; a striking example is shown in Figure 8d.

**Figure 8:**
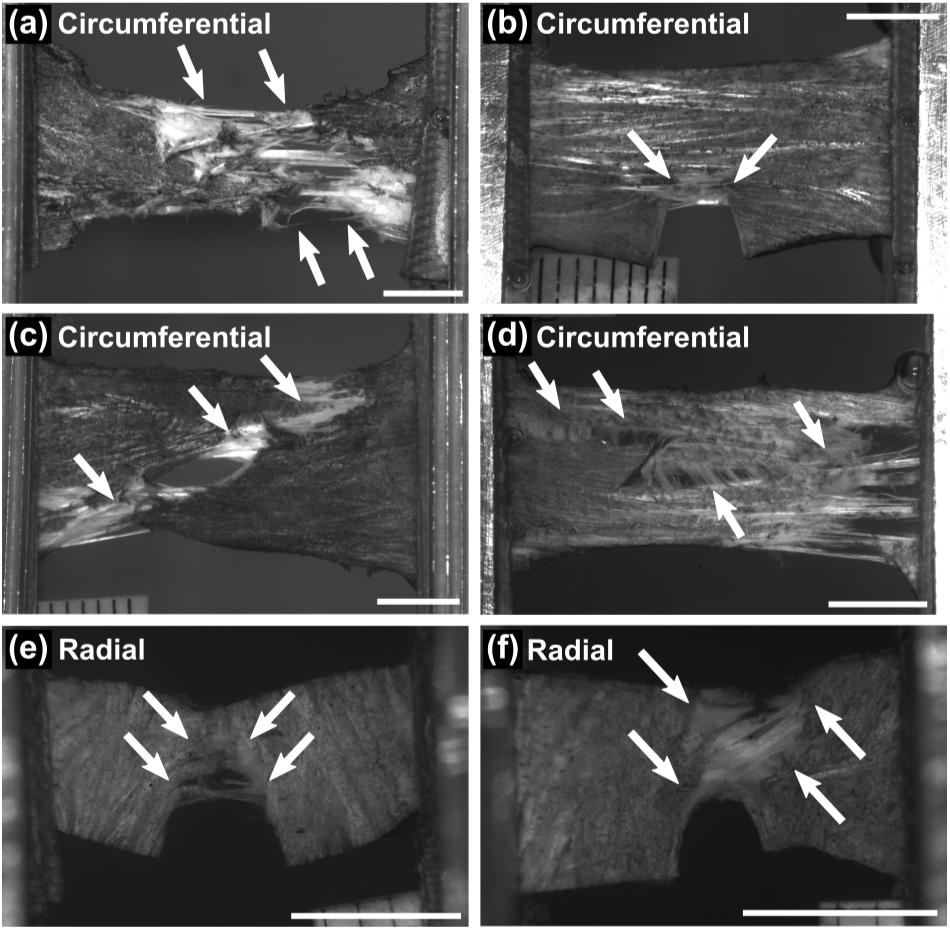
Examples of cracked specimen rupture morphology. (a) Circumferential control specimen, showing a widespread rupture zone and interdigitating fiber pullout that is typical of circumferential specimen rupture. (b) Circumferential edge crack specimen, showing early rupture at the crack tip combined with crack blunting by inter-fascicle shear. (c) Circumferential center crack specimen, showing the typical outcome of independent rupture sites merging with the crack. (d) Circumferential center crack specimen, showing zigzagging of a rupture across the crack and along fascicle interfaces. (e) Radial edge crack specimen, showing crack growth. (f) Radial edge crack specimen, showing simultaneous rupture across its entire width. (all) Scale bar = 5 mm.

Radial specimens, unlike circumferential specimens, sometimes displayed crack growth (incremental, local rupture growing from the crack tip) (Figure 8e). In 9/17 radial tests, the crack tip-associated rupture grew incrementally such that total rupture occurred by crack growth. In 5 of these 9 tests a separate site of rupture developed away from the crack in parallel with the crack growth. Crack growth did not begin until strain approximately reached ε_u_ ± 0.2ε_u_. The 8/17 radial tests with no sign of crack growth ruptured such that the site of rupture touched the crack in 3 tests (Figure 8f) and was away from the crack in 5 tests. Most (16/17) radial tests had some degree of local rupture at the crack tip even if this did not translate into crack growth. Regardless of the type of rupture, the rupture site was bridged by small fibers (Figures 8e,f). Although in the circumferential specimens the fascicle interfaces appeared to facilitate the formation of shear bands, in the radial specimens they appeared to facilitate incremental crack growth.

The effect of a crack on the type of rupture (midsubstance, mixed, and grip line) was examined by comparing counts of rupture types between cracked specimen groups and their controls (Table 2). Circumferential edge crack specimens had significantly more mixed ruptures, fewer grip line ruptures, and slightly fewer midsubstance ruptures (relative to the total number) than controls (p = 0.04, chi-squared test). Rupture type counts for circumferential center crack specimens were not significantly different from controls. Radial edge crack specimens had significantly more midsubstance and mixed ruptures and fewer grip line ruptures than their controls (p = 0.04, chi-squared test).

## 4 Discussion

### 4.1 Summary of key findings

This study’s objectives were to determine whether a short crack in meniscus tissue reduces the meniscus’ ultimate stress in tension and hence requires failure analysis by fracture mechanics methods and, furthermore, to determine whether a crack alters the strain field by creating a near-tip strain concentration and an unloaded region in the cut material on either side of the crack. No reduction in ultimate stress in cracked specimens was detected relative to controls (Figure 3), indicating that limit stress analysis is appropriate for failure analysis, and that fracture mechanics analysis is not required, in the tested conditions. Since the study was adequately powered to detect a meaningful (~ 1 s.d.) decrease in ultimate stress, these results are robust evidence for preservation of ultimate stress in the presence of a short crack. Regarding strain concentrations, greater tissue strain was observed near the crack tip (Figure 5), implying the existence of a corresponding near-tip stress concentration. Consistent with the presence of near-tip strain concentrations, the region containing material cut by the crack (the cut ROI) had reduced strain, implying that it was partially unloaded (Figure 7). Although the alterations in the strain field did not measurably influence the meniscus’ rupture in terms of the overall stress–strain curve, the near-tip stress concentration does pose a risk of sub-rupture local tissue damage.

### 4.2 Significance of preservation of ultimate stress

The preservation of ultimate stress in cracked specimens and the tendency for ruptures to occur simultaneously through the entire cross-section implies a high degree of resistance to crack growth (fracture toughness). Although half of radial specimens exhibited crack growth, this occurred at or past the ultimate stress point, by which time rupture was well underway by other mechanisms. There was no evidence of unstable crack growth. This crack growth may therefore be considered incidental, similar to the final fracture of a highly tough and ductile test piece (Broberg 1999). High fracture toughness is desirable for meniscus because the maintenance of strength, despite the presence of a crack, lessens the risk of further failure and loss of function. Note that a crack will still reduce the cross-sectional area and, in proportion, the total force that the meniscus can support. However, the present study’s results indicate that short (~ 3 mm) tears in single-cycle loading are unlikely to accelerate failure by crack growth.

Longer cracks, in principle, produce stronger stress concentrations. In vivo meniscus tears that are considered to definitely require clinical treatment are 5 mm to 25 mm long (Stärke et al. 2009; Roeddecker et al. 1994; Lento and Akuthota 2000; Seil et al. 2009). These tears may exceed the critical crack length for fracture mechanics-dominant rupture (i.e., fracture) (Taylor 2007). However, the apparently low risk associated with a short (natural) tear is still beneficial because it should delay the growth of the tear to a critical size. Attempts to replace meniscus tissue with engineered materials should consider replication of the meniscus’ fracture toughness as a design requirement.

This assessment of low risk for a short meniscus tear has caveats, as it is based on excised tissue tests with test conditions that were, by necessity, artificial. Specimen size and hence crack length was small, only edge and center cracks were examined, specimens were cut only from the center of the meniscus, specimens were loaded only in circumferential and radial uniaxial tension, cutting specimens may cause loss of fiber integrity, the loading rate was relatively slow (0.5 mm/s), and unlike the natural enthesis the tensile grips work by compression and friction rather than by individually anchoring each fiber. In vivo, tears come in a wide variety of geometries and the meniscus is subjected to surface contact and multiaxial, heterogeneous stress (Kawamura et al. 2003; Fithian et al. 1990; Mononen et al. 2013; Aspden 1985; Spilker et al. 1992; Párraga Quiroga et al. 2014; Atmaca et al. 2013; Freutel et al. 2014). Considerable work remains to develop laboratory test methods that reproduce the wide variety of loading conditions under which the meniscus must function in vivo, and to quantify damage and failure in those conditions. Pending development of more advanced methods, however, the present results support a low risk of total rupture due to short tears.

### 4.3 Significance of crack tip strain concentrations

Although cracks did not decrease the meniscus specimens’ ultimate stress, they did create regions of greater strain near their tips. These strain concentrations were substantial. Depending on the test scenario and stress level, mean longitudinal strain was increased by 20% to 50% and mean shear strain magnitude was increased by 20% to 80% (Figure 5). Groups with more complete unloading of the cut ROI (circumferential and radial edge crack specimens) also had greater near-tip E_xx_ concentrations, consistent with load redistribution around the crack being the cause of the strain concentrations. This near-tip overstrain has potential to cause local damage and failure under normal load levels. Damage is generally reported to begin at 0.05–0.08 strain in tendon and ligament (Provenzano et al. 2002; Duenwald-Kuehl et al. 2012; Sverdlik and Lanir 2002; Zitnay et al. 2017), with one study showing a damage threshold ≈ 0.02 strain (Lee et al. 2017), but thresholds for damage or permanent deformation are not known for meniscus. If the meniscus is similar to tendon and ligament, the stress threshold at which a crack makes meniscus tissue vulnerable to damage would be 25–50% of its tensile strength; that is, 9–17 MPa. At these stress levels, crack blunting via inter-fascicle shear was prevalent in circumferential edge and center crack specimens and rupture from the crack tip along fascicle boundaries was prevalent in circumferential edge crack specimens. These findings indicate that the near-tip shear concentrations facilitated sliding and rupture at the fascicle interfaces. Furthermore, the existence of zones of weakness near the tips of natural meniscus tears suggests that damage due to strain concentrations also occurs in vivo (Roeddecker et al. 1994). Local strain concentrations created by cracks in meniscus tissue therefore pose a high risk of causing local, sub-rupture damage.

### 4.4 Combined interpretation of preservation of ultimate stress and strain concentrations

The preservation of ultimate stress and general lack of crack growth in cracked meniscus tissue stands in contrast to the high risk of local damage due to strain concentrations near the crack tip. These risks may have an inherent trade-off. Fracture toughness represents the energy required for crack growth. Local energy dissipation around the crack tip (in the process region) increases fracture toughness (Taylor et al. 2012; Janssen et al. 2002). The observed near-tip zone of local overstrain and probable damage may function as a dissipative zone, with the function of increasing toughness and limiting crack growth. Inter-fascicle shear, which was especially strong in circumferential specimens, may increase the meniscus’ toughness by absorbing energy through fascicle sliding. Inter-fascicle sliding also deflects the path of rupture along fascicle boundaries (e.g., Figure 8d), which due to the circumferential specimens’ geometry promotes rupture by processes other than mode I crack opening and crack growth. In radial specimens, the parallel alignment of the fascicles and the crack prevented fascicle sliding from acting as a crack deflection mechanism, which is probably why half of radial specimens exhibited crack growth, albeit still without measurable impact on ultimate stress. The roles of local damage, plasticity, and crack deflection as toughening mechanisms in the meniscus warrant further investigation.

Strain concentrations near the tip of a meniscus tear, when subjected to long-term repeated loading, may cause incremental accumulation of damage and failure. In the circumferential tests, shear strain concentrations extended 5–10 mm from the crack tip along fascicle boundaries and were halted by the grip line. In vivo they would not be restricted in this manner and hence could provide a path for progressive growth of circumferential, bucket handle, vertical, or horizontal tears (Aspden 1985; Fithian et al. 1990; Kelly et al. 1990; Smillie 1978). A strain concentration may also kill or damage cells even in a single loading cycle. In ligament, cell death has been shown to follow a linear no-threshold relationship with applied strain (Provenzano et al. 2002). A meniscus tear is also likely to cause regional unloading as observed here for artificial cracks (Figure 7), which may lead to pathology due to cells’ dependence on load to maintain homeostasis (Natsu-Ume et al. 2005; Frank et al. 2004; Gupta et al. 2008). Altered cell loading and incremental accumulation of tissue and cell damage may mediate long-term development of in vivo meniscus pathology.

### 4.5 Increased tangent modulus and yield stress in edge crack circumferential tests

The observation of greater tangent modulus and yield stress in edge crack circumferential tests compared to controls (Figure 3) was unexpected and its cause is indeterminate. One possibility is that it occurred by chance; i.e., that it is a type I statistical error. A second possibility is that the inner margin of a circumferential specimen has an effective stiffness less than the outer margin. If so, the edge crack, by eliminating the low-stiffness inner margin from the intact cross-section, would cause the average modulus to increase. A lesser effective stiffness in the inner margin implies that the ultimate stress for circumferential edge crack specimens should have also increased; this is neither confirmed nor ruled out by the data (p = 0.07, post-hoc two-tailed t-test). A lesser effective stiffness in the inner margin of circumferential specimens, if present, could have been caused by differences between the inner and outer margins in either (a) material properties or (b) fascicle loading.

The meniscus’ material properties, specifically tissue composition and stiffness, have been observed to vary with radial position in the meniscus (Cheung 1987; Kelly et al. 1990; Nakano et al. 1986; Ghosh et al. 1983; Fithian et al. 1990). However, these comparisons were of the outer, central, and inner thirds, and the specimens in the present study were all cut from the central third. Electron microscopy and histology show homogeneous composition in the central third (Petersen and Tillmann 1998; Andrews et al. 2014). There is no prior evidence that the inner margin of specimens in the present study should have significantly different material properties from the rest of the specimen. The observation of greater tangent modulus and yield stress in edge crack circumferential specimens is therefore unlikely to have been caused by variation in material properties with radial position.

A reduction in fascicle loading within the inner margin of circumferential specimens could have been caused by the specimens’ geometry. The meniscus’ fascicles are aligned in arcs, but the specimens’ edges were straight. A portion of the inner margin’s fascicles were therefore discontinuous with the gripped region and were loaded indirectly by inter-fascicle shear. The fascicle interfaces are apparently compliant and shear readily (Figures 6 and 8b), so load transfer to the discontinuous fascicles is likely to be slight. This effect has been quantified in annulus fibrosus and tendon (Skaggs et al. 1994; Szczesny et al. 2015). The inner margin’s fascicles may therefore be relatively unloaded, contributing little to the stiffness of the specimen. Unloading of inner margin fascicles would artificially decrease the average modulus in circumferential control and center crack specimens, which have an intact inner edge, but not in edge crack specimens because the crack eliminates these inner margin fascicles from the load-bearing cross-sectional area.

The potential for fascicle unloading due to the mismatch between straight sides of conventional tensile test specimens and their arced fascicle geometry deserves further investigation. Interactions between specimen geometry and fiber continuity have been previously shown to alter the meniscus’ apparent tensile properties (Peloquin et al. 2016), prompting the use of expanded grip regions. However, the present tangent modulus and yield stress results suggest that the meniscus’ arced fascicle architecture remains a challenge. In vivo, the meniscus’ entheses likely provide much more efficacious fiber loading than is currently possible in laboratory tests.

### 4.6 In vivo tear growth and future directions

The general lack of crack growth observed in this study, especially in the circumferential tension tests, leaves the mechanisms by which meniscus tears grow in vivo uncertain. In particular, it would be very useful to resolve uncertainty regarding the duration for which a short in vivo meniscus tear can be expected to remain short and hence low-risk. The observation of crack growth along fascicle boundaries in half of the radial specimens suggests that meniscus microstructure is important, although many in vivo tears do cross fascicles. High cycle loading, as has recently been undertaken for crack-free meniscus (Creechley et al. 2017), is a promising approach. Loading rate is another potentially important factor. Meniscus tears often occur in the context of sports, implying involvement of high-rate loading (Drosos and Pozo 2004; Feucht et al. 2015), and ex vivo impact testing has produced meniscus tears (Isaac et al. 2010; Isaac et al. 2008). It is possible that a high-rate loading cycle (as would be expected from an impact) can produce a long (5–25 mm) tear of clinical concern immediately, without slow growth from a low-risk short tear. Although the focus of this study was on tensile loading, examining the behavior of meniscus cracks under shear is also a promising direction for soft tissue (Haslach et al. 2018; Taylor et al. 2012). The central task for future experiments is to produce physiologic growth of cracks under controlled laboratory conditions so that the mechanisms and time course of this pathology can be quantified.

## 5 Data accessibility statement

The underlying data for all results is available, hosted on the Open Science Framework, under a CC-BY license (Peloquin 2018).

## 7 Acknowledgments

Julia Pezick and Pranita Muralidhar contributed a great deal to this work by performing many of the mechanical tests.

## 8 Supporting information captions

**Figure S1.**
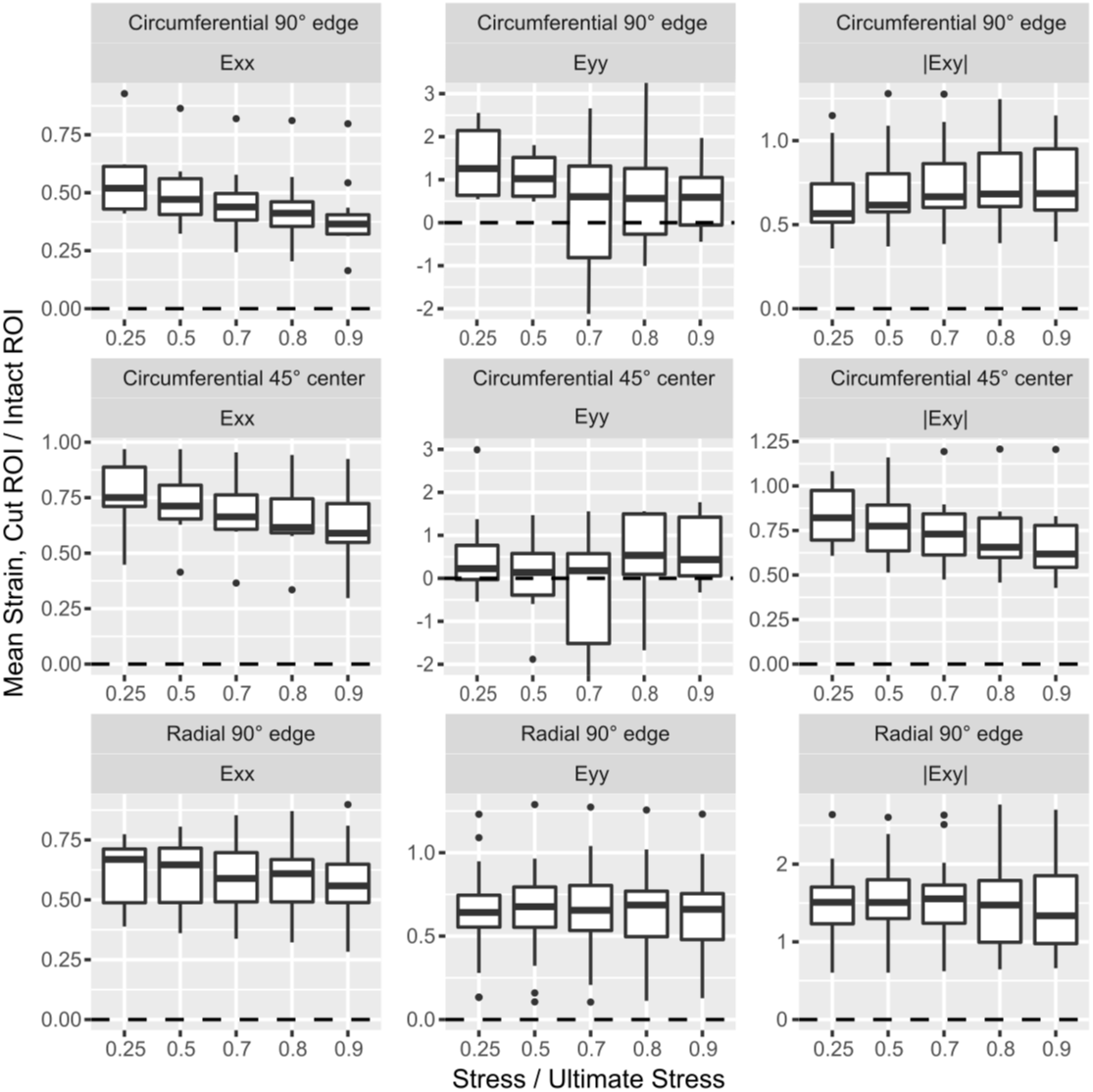
Ratios between mean strain in cut and intact ROIs. The plots of E_yy_ ratios for the circumferential edge and center crack groups are cropped to exclude 3 specimens each for which intact strain ≈ 0, producing a ratio ≈ ∞. All plotted quantiles remain based on the full dataset. The boxplots are in Tukey’s style.

**Table S1. Complete stress-strain curve statistics for all groups.** See supplemental files.

**Table S2. Complete strain field statistics for near-tip and away-from-tip ROIs for cracked groups.** See supplemental files.

